# Short term treatment with a cocktail of rapamycin, acarbose and phenylbutyrate slows aging in mice

**DOI:** 10.1101/2021.10.21.465380

**Authors:** Zhou Jiang, Juan Wang, Denise Imai, Tim Snider, Ruby Mangalindan, John Morton, Lida Zhu, Adam B. Salmon, Jackson Wezeman, Jenna Klug, Jiayi Hu, Vinal Menon, Nicholas Marka, Laura Neidernhofer, Warren Ladiges

## Abstract

Pharmaceutical intervention of aging requires targeting multiple pathways, thus there is rationale to test combinations of drugs each targeting different but overlapping processes. In order to determine if combining drugs previously shown to improve lifespan would have greater impact than any individuyal drug, a diet containing rapamycin at 14 ppm, acarbose at 1000 ppm, and phenylbutyrate at 1000 ppm was fed to 20-month-old C57BL/6 and HET3 4-way cross mice of both sexes for three months. Mice fed the cocktail diet showed a strain and gender-dependent phenotype consistent with healthy aging including decreased body fat and blood glucose, improved cognition, and increased grip strength and walking ability compared to mice fed individual drug or control diets. A cocktail diet containing ½ dosing of each compound was overall less effective than the full dose. The composite age-related lesion score of heart, lungs, liver and kidney was decreased in mice fed the cocktail diet compared to mice fed individual drug or control diets suggesting an interactive advantage of the three drugs. Senescence and inflammatory cytokine levels in kidneys from mice fed the cocktail diet were lower than in kidneys from mice fed control diet, and consistent with low expression levels in kidneys from young untreated mice, suggesting the cocktail diet delayed aging partly by senolytic and anti-inflammatory effects.

## Introduction

The basic biology of aging affects the functional performance of all organs in the body. The field of geroscience aims to understand, at the cellular and molecular level, the interconnections between aging and disease/disabilities, with a focus on understanding the mechanisms by which aging contributes to most chronic diseases as a major risk factor^1^. The geroscience concept posits that manipulation of aging will simultaneously delay the appearance or severity of multiple chronic conditions and diseases because they share the same underlying major risk factor: aging and the multiple processes involved in aging^2,3^.

Recent progress in the field of aging biology has allowed researchers to develop robust behavioral, genetic and pharmacological approaches to expand the lifespan of multiple species. Importantly, interventions that extend lifespan often result in improvements in multiple aspects of health span, resulting in significant delays in the appearance of pathology and frailty^4^. We selected a drug cocktail of rapamycin (Rap), acarbose (Acb), and phenylbutyrate (Pba) to test this concept in aging mice. The rationale for the drug cocktail is based on validated anti-aging effects of the individual drugs each targeting different but overlapping processes of aging.

Rap is an antibiotic clinically approved for treating organ transplant patients^5,6^ and neoplastic conditions. It is orally bioavailable and readily crosses the blood brain barrier. Rap inhibits mTOR signaling which is plays a significant role in integrating signals from growth factors and nutrients to control protein synthesis. The effect of down regulating mTOR on aging was confirmed by the NIA Intervention Testing Program showing that rapamycin extended lifespan in mice^7^. It has also been shown to delay age-associated heart disease and other pathologies in mice^8,9,10^. Transient Rap treatment has been shown to increase lifespan and healthspan in middle age mice^11^. Rap and derivatives are currently being tested in clinical trials for numerous chronic disease conditions.

Acb is a popular type 2 diabetes medication used for glucoregulatory control^12^. Considering that one of the hallmarks of caloric restriction (CR) is improved glucose homeostasis and insulin sensitivity^13^, the similarities between CR and Acb treatment underscore the potential metabolic responses that are present in conditions of improved aging. Like CR, Acb reduces body weight and body fat, improves glucose dysregulation associated with aging, and increases mouse lifespan^14^.

Pba is clinically approved as an ammonia scavenger for urea cycle disorders in children. It is a derivative of the short chain fatty acid butyrate, which occurs naturally in the gut. It is orally bioavailable and can be detected in the blood stream within 15 minutes of oral administration, and organs within an hour. We have shown that Pba can enhance physical and cognitive performance in aging mice^15^. Pba is currently being tested in both preclinical and clinical studies for several age-related conditions.

These three drugs each target different aging processes. Rap targets autophagy and vascular deficits by downregulating mTOR signaling. Acb indirectly targets insulin signaling, mitochondrial dysfunction and oxidative stress by blocking breakdown of complex carbohydrate in the small intestine so that less glucose is available for systemic absorption. Pba targets dysfunctional proteostasis through the endoplasmic reticulum stress response and targets epigenetic function by inhibiting histone deacetylation resulting in upregulation of genes including anti-inflammatory genes. We posit that our cocktail will be successful in delaying aging because it targets more aging processes than any individual drug.

## Methods

### Animals and drug treatment

C57BL/6 (B6) and HET3 4-way cross (H3) mice of both genders were used for this study. B6 mice were received from NIA at the age of 20 months. H3 mice were provided by the Nathan Shock Center at the University of Texas Health San Antonio. Mice were acclimated two weeks before randomly assigned to different diet treatment. Mice were kept under 12 hours day/night monitoring and specific pathogen free condition and with free access to food and water. Mice were started on the study at 20 months of age and continued for three months, when the study ended at 23 months of age. Each cohort had 20 mice, 10 males and 10 females. All experiments were conducted in accordance with approval by the University of Washington IACUC.

Drugs were delivered orally in the diet, which was prepared by TestDiet, Inc, a division of Purina Mills. Five different drugs or drug combinations were prepared plus a non-medicated control diet. Doses for Rap, Acb and Pba were those that enhanced health span and lifespan as reported for mice^15,16,17^. Microencapsulated Rap was obtained from Southwest Research Institute (San Antonio, TX) and mixed at a concentration of 14 ppm. Acb was obtained from Spectrum Chemical Mfg Corp., Gardena, CA and mixed at a concentration of 1000 ppm. Pba was obtained from Triple Crown America, Inc, Perkasie, PA under the commercial name of sodium triButyrate, and mixed in the feed at a concentration of 1000 ppm.

Mice were weighed weekly. Lean and fat mass were recorded monthly through a calibrated QMR (EchoMRI) imaging system. Food consumption was monitored monthly over a three-day period wherein food weight difference was compared between 0 and 72 hours. The amount of food consumed by each mouse was estimated by the food weight difference divided by number of mice in the cage. Blood glucose levels were measured monthly using tail blood with a glucometer (CONTOUR®NEXT EZ meter).

### Performance assays

Rotarod is a test for coordinated walking ability, and was performed as described^18^ using a rotarod apparatus (Rotamax 4/8, Columbus Instruments, Inc.) that allowed mice to walk on a rotating rod with constantly increasing rotation speed. Mice were placed in the lanes of the rotarod with initial rod speed at 0 RPM. The speed was progressively increased by 0.1 RPM/sec (0 to 40 RPM over 5 minutes) or until all mice were dislodged as determined by an infrared sensor. The time in seconds was recorded for three trails with half hour resting time in between each trail.

Hand grip is a universal measurement used to assess physical competency in older adults as an indicator of frailty. The grip strength test in mice is similar to the hand grip test for people in that it assesses the ability to grip a device with the front paws^19,20^. Mice were positioned horizontally from a grip bar (Columbus Instruments, Inc.) and pulled back slowly and steadily until they released their grip. The test was repeated five times and peak force for the forelimb paws was recorded. Grip strength force was normalized by body weight measured on the testing date, so that the peak force was expressed relative to body weight.

Cognitive function was compared between mouse cohorts using a spatial navigation task (Box maze), a learning paradigm which reflects working memory and cognitive flexibility^21^. In this task, mice were introduced into a bright special box with seven blocked exit and one escape exit leading to a dark standard housing cage. Four consecutive trials were given to each mouse with 120 second maximum maze exploiting time and 30 seconds resting period in between each trial.

### Quantitative Real-Time PCR

Tissues were snap frozen to −80 C immediately after dissection. Homogenized frozen tissues were concentrated for RNA extraction with measured purity and relative quantity. 5 µg of total RNA was used for reverse transcription synthesis of CDNA. SYBER Green was chosen for dsDNA binding to quantify PCR product.

### Geropathology

Mice were humanely euthanized according to USDA guidelines. Heart, lung, liver and kidney were fixed in 10% buffered formalin for 48 hours then blocked in paraffin wax through University of Washington Department of Comparative Medicine’s Histology Lab. Sections of 4 µm were stained with hematoxylin and eosin for blind geropathology grading by two board certified veterinary pathologists (D Imai and T Snider) according to published guidelines^22^. The geropathology scores were tabulated into a composite lesion score (CLS) of four major organs-heart, lungs, liver and kidney, and recorded as a single value for each animal.

### Data Analysis

Two-tailed student’s t-test was used to compare between results from each treatment cohort. Chi-square tests were used to determine the association between categorical outcomes and treatment cohorts. For geropathology, scoring data was the average lesion score from two pathologists analyzed by cohort using the two-tailed student’s t-test. Mean values with standard error bars (SEM) are presented by group in the figures. Quantitative interval outcomes collected from both pathologists were normalized for central tendency analysis. A mixed-effects model with random intercept was applied to investigate the qPCR results of senescence associated factor expression^23^. The marginal composite scores were tested through log-transformed relative expression levels. Correlation analysis between treatment cohorts, strains and lesion scores was done by one- and two-way ANOVA. All statistical tests were conducted at 0.05 significance level.

## Results

### Mice fed a diet containing a cocktail of rapamycin, acarbose, and phenylbutyrate for three months showed strain and gender-dependent biological changes, superior performance, and decreased geropathology compared to mice fed a non-medicated diet

The percent change for body weight, body fat mass, lean muscle mass, blood glucose, and food consumption from average baseline values is shown in Figure 1A. Changes in body fat mass (BFM) were highly sensitive to the drug cocktail. In B6 mice, BFM decreased by an average of 36 percent in females and 33 percent in males compared to 10 percent in females and no change in males on the control diet. The effect was just as robust in H3 mice with females showing a 37 percent decrease and males showing a 45 percent decrease.

**Figure 1.**
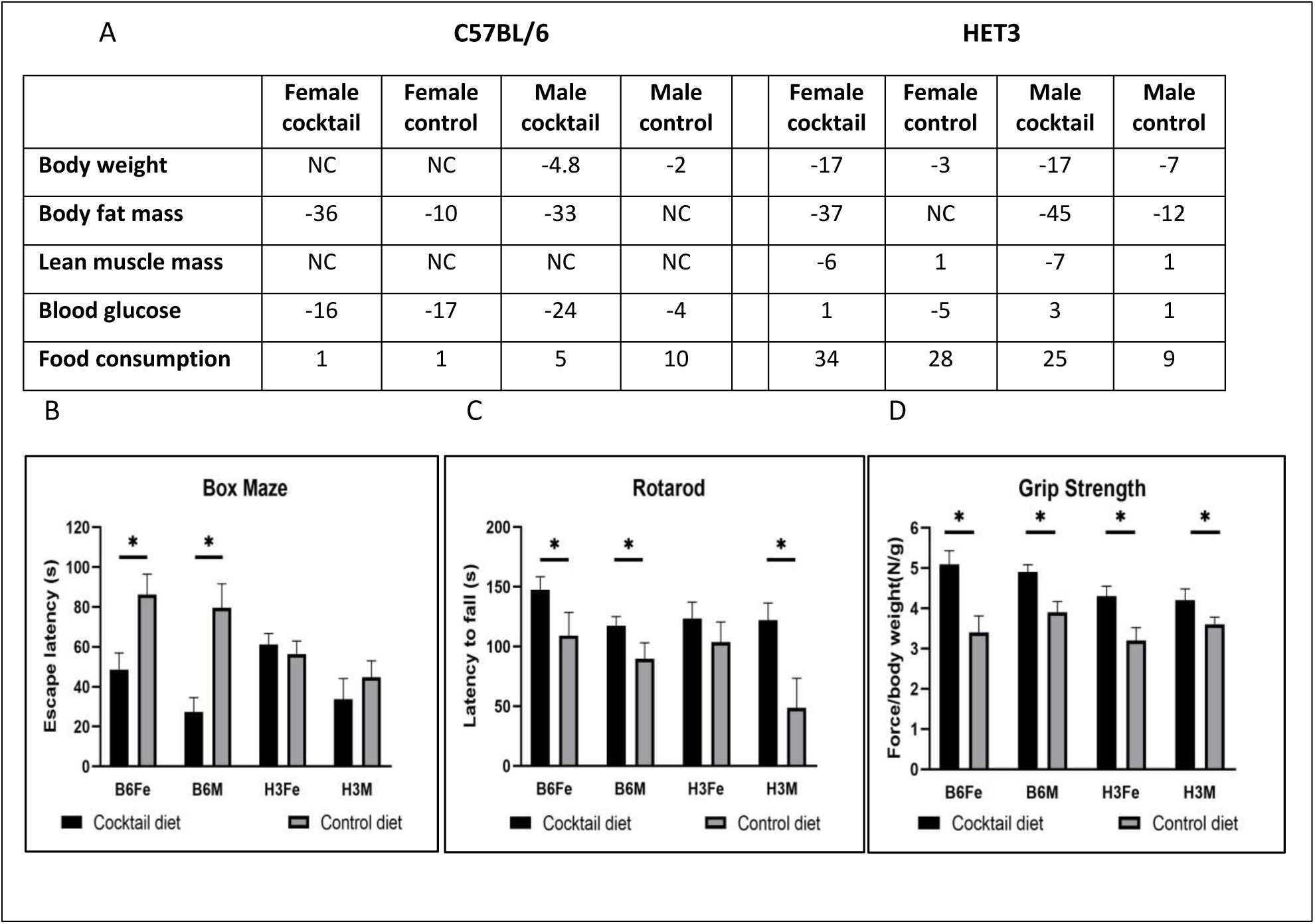
**A**. C57BL/6 and HET3 mice showed changes in biological values (designated as percent change) after three months on a drug cocktail (rapamycin, acarbose, and phenylbutyrate) diet starting at 20 months of age compared to mice fed a control diet. **B)** Performance tests were conducted in mice at 23 months of age, three months after being on a diet containing the drug cocktail or placebo. Values for the box maze are standardized times in seconds to find the escape hole in trial 3. **C)** Values for the rotorod are standardized times in seconds staying on the rotating rod. **D)** Values for grip strength are standardized force in neutons per body weight measuring the ability to maintain forepaw grip from a parallel meter bar. N = 9-10, *p≤0.05.

Both female and male B6 mice exhibited age related cognitive impairment which was robustly attenuated in mice fed the cocktail diet (Fig 1B). H3 mice were not as cognitively impaired but H3 males treated with the cocktail did show improved cognition. B6 females and H3 males treated with the cocktail showed enhanced performance on the rotarod, while B6 males and H3 females did not (Figure 1C). Both genders in both strains treated with the cocktail showed increased grip strength compared to cohorts treated with placebo (Fig 1D).

We wanted to see if there was any strain and gender differences in how the cocktail affected age related lesions in H3 mice, a commonly used strain in aging studies, compared to B6 mice. Age related lesion severity was decreased in both female and male B6 mice and H3 female mice fed the cocktail diet (Figure 2). H3 males fed the cocktail diet showed a lesion severity similar to mice fed the control diet suggesting there were gender and strain differences in the cocktail response to age-related lesions.

**Figure 2.**
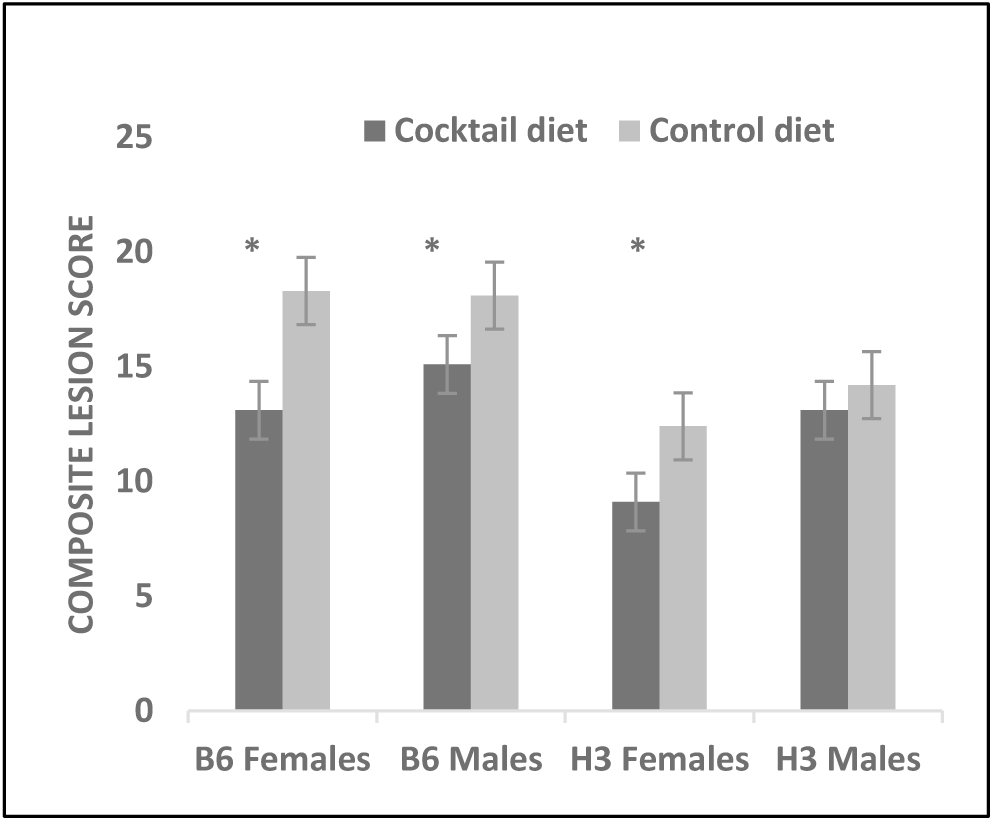
The cocktail diet was more effective in decreasing lesion severity in C57BL/6 mice than in HET3 mice. The composite lesion severity decreased in C57BL/6 females and males and HET3 females but not males fed a drug cocktail diet for three months starting at 20 months of age compared to composite lesion severity mice fed the control diet. *p≤0.05. N = 10-12/cohort.

### A half dose of the drug cocktail diet fed to C57BL/6 mice was overall less effective than the full dose cocktail

We wanted to see if a lower dose of each of the three drugs administered as a cocktail would show the same effectiveness as the full dose. Both males and females consistently performed better in the spatial navigation task (Box maze) when fed the full cocktail diet or the half cocktail diet compared to the control diet (Figure 3B). However, the full cocktail diet was more effective than the half cocktail diet or control diet in females performing the rotarod task and in males performing the grip strength task (Figure 3C and D).

**Figure 3.**
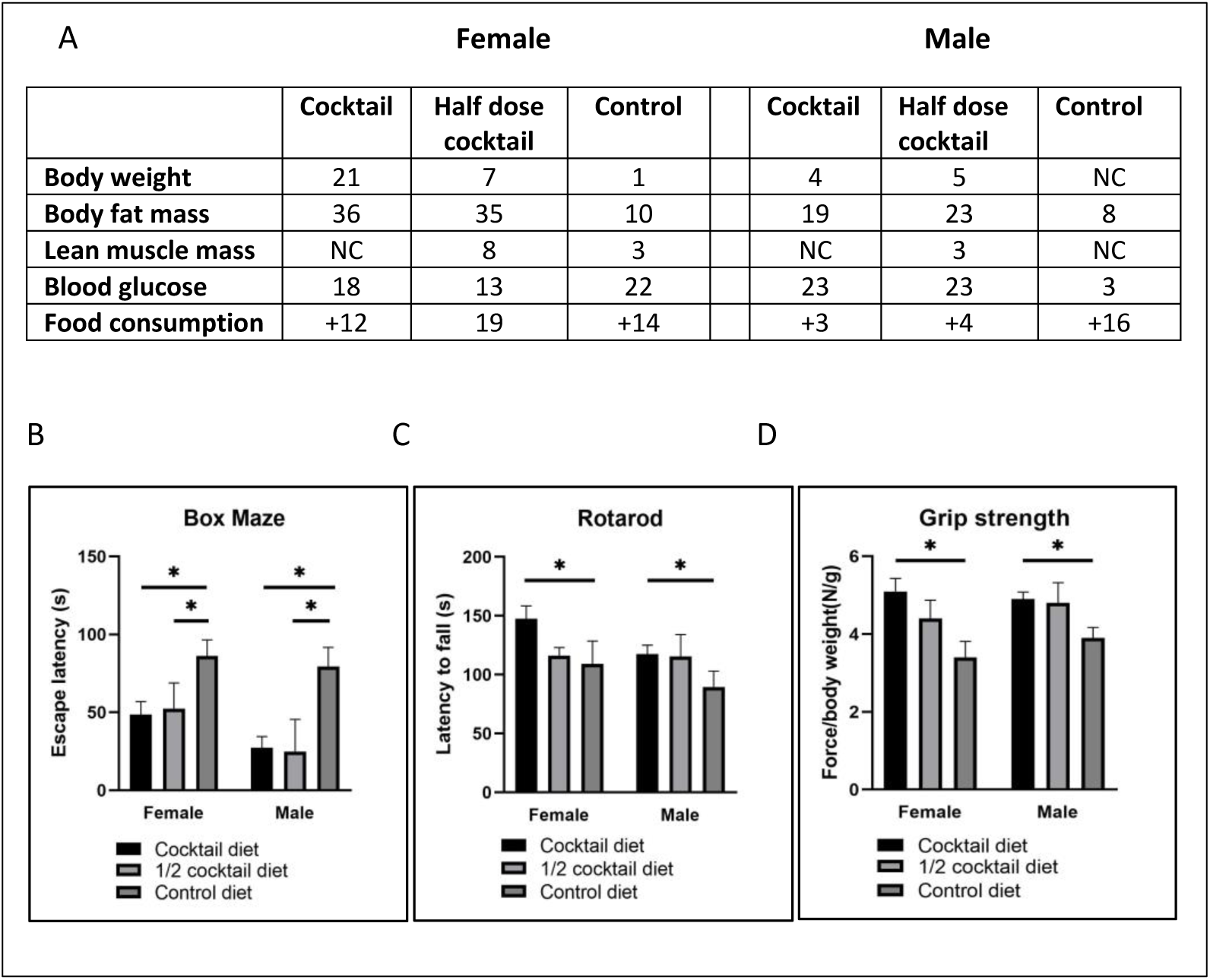
C57BL/6 mice fed the half dose cocktail diet had similar biological values but only partial performance test results compared to mice fed the full dose cocktail diet. **A**. Mice receiving the half cocktail diet had similar biological values compared to the full cocktail diet with a few exceptions. **B**. Both males and females fed full or half cocktail diets found the escape hole more quickly than mice fed the control diet. **C**. Male mice fed the half cocktail diet outperformed female mice in the rotarod in a manner similar to mice fed the full cocktail diet. **D**. Female mice fed the half cocktail diet outperformed male mice in grip strength in a manner similar to mice fed the full cocktail diet. N = 9-10, p≤0.05.

The half dose cocktail diet was not effective in decreasing the severity of age-related lesions in either B6 females or males compared to the full dose drug diet (Figure 4).

**Figure 4.**
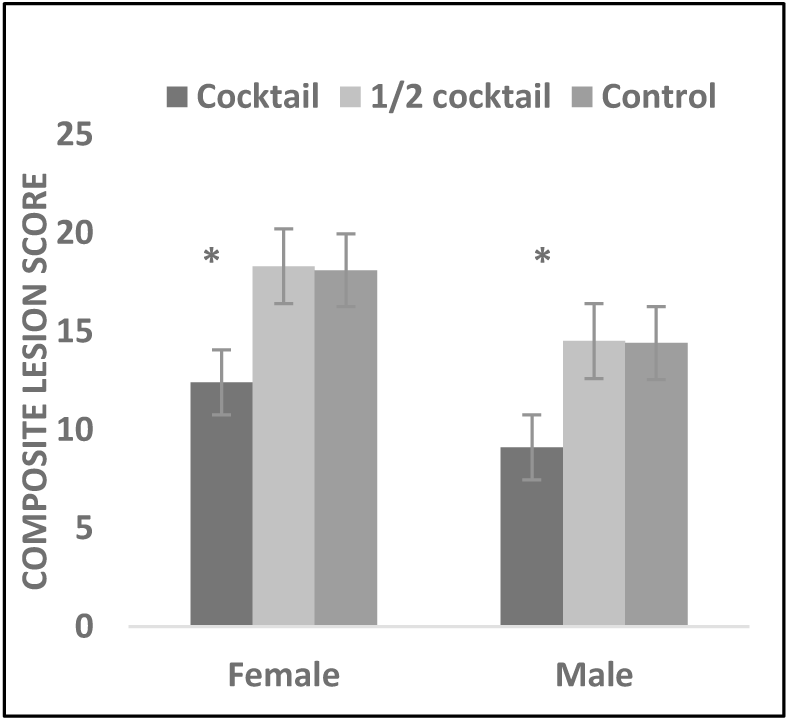
The cocktail diet was more effective in decreasing lesion severity compared to the half dose cocktail diet in both C57BL/6 females and males. *p≤0.05, N=10-12/cohort.

### The drug cocktail diet was more effective than each individual drug diet in enhancing healthy aging

We next wanted to see what effect each individual drug had on aging parameters over the course of three months compared to the combination of these drugs. There was considerable variation in biological values among all of the cohorts at the end of three months (Figure 5A). Acb increased food consumption and decreased body fat mass in both females and males, which was partly reflected in the cocktail cohort. Both male and female mice fed the Rap diet performed better in the spatial navigation task than mice fed Acb and Pba diets and equaled performance of mice fed the cocktail diet (Figure 5B). In general, both male and female mice fed the cocktail diet outperformed mice fed individual drug diets in rotarod and grip strength tasks (Figure 5C and D).

**Figure 5.**
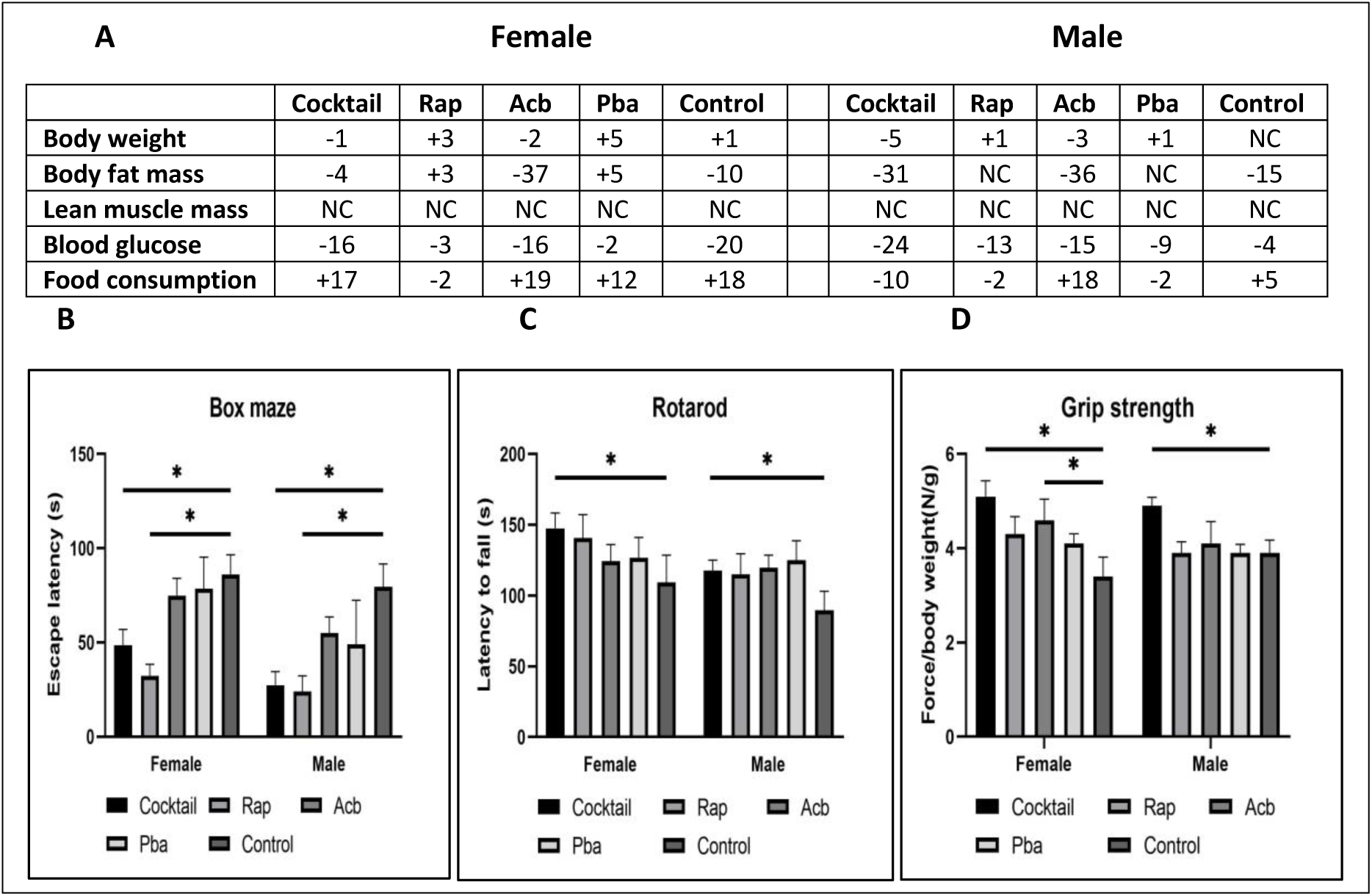
In general, B6 mice fed the cocktail diet for 3 months showed decreased body weight, fat mass and blood glucose and increased performance compared to mice fed each individual drug or control diet. **A**. Biological values; **B**. Box maze values; **C**. Rotarod values**; D**. Grip strength values. N = 9-10, p≤0.05.

There was a decrease in lesion severity in both males and females fed the cocktail diet (Figure 6). The individual Rap, Acb or Pba diets were no more effective than the control diet in decreasing composite lesion severity suggesting an interactive advantage of the three drugs adminstered together over the same time.

**Figure 6.**
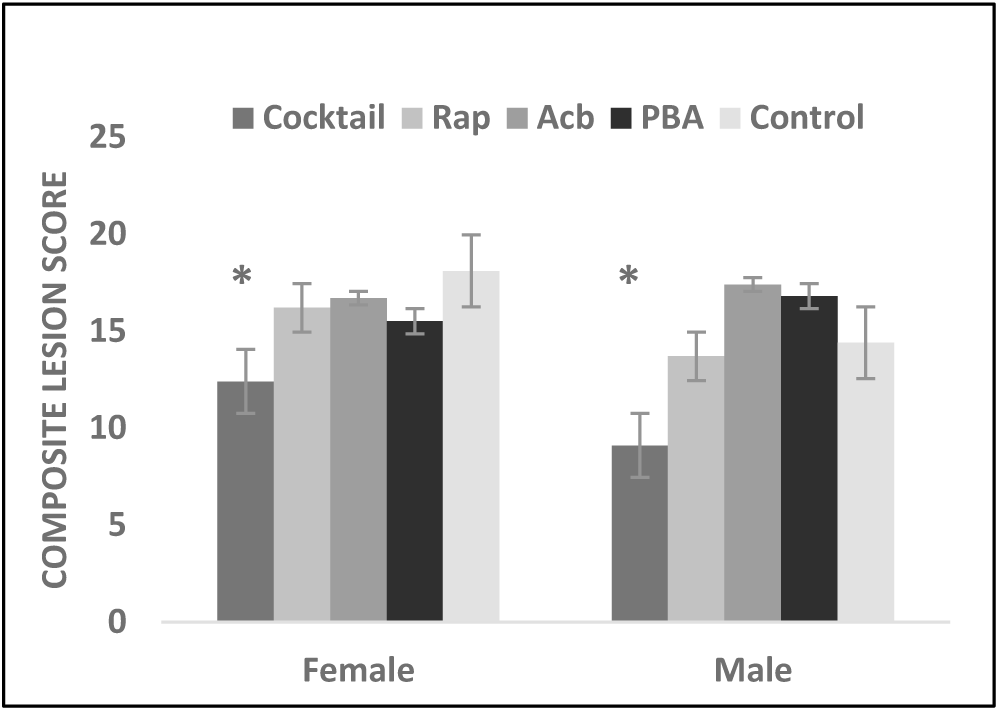
The cocktail diet decreased the severity of age-related lesions while the individual drug diets did not. C57BL/6 mice fed the cocktail diet for 3 months had a greater decrease in severity of age-related lesions compared to mice fed each individual drug diet or control diet. *p≤0.05, N = 10-12/cohort.

### The drug cocktail lowered levels of senescence and inflammatory cytokines in the kidney

RNA message levels for TNFa, IL6, and IL1b in the kidneys from cocktail treated mice were assessed by rtPCR and shown to be significantly lower than the kidneys from control treated mice (Figure 7). The lower expression levels were consistent with low expression levels in the kidneys from young untreated mice. Similar findings were found for p16 expression levels.

**Figure 7.**
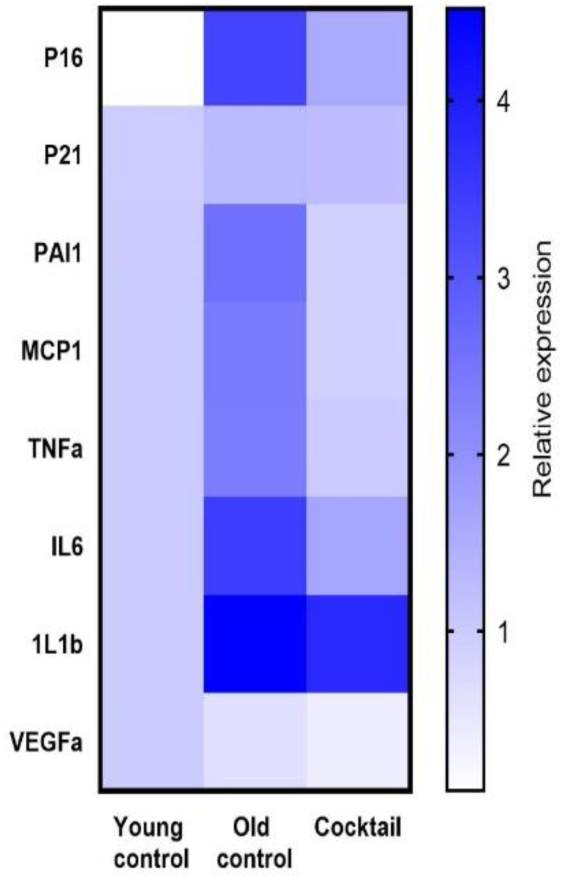
Kidneys from 23-month-old C57BL/6 mice fed the cocktail diet for 3 months showed a general decrease in senescence and inflammatory associated proteins by qPCR. Based on the mixed-effect model, which accounts for individual and composite relative expression values, the cocktail diet group showed significantly more senolytic effect than the control diet group (p<0.001) and more in line with expression in young mice.

## Discussion

The drug cocktail of rapamycin (Rap), acarbose (Acb), and phenylbutyrate (Pba) was studied to test the concept that interventions that extend lifespan in mice will result in improvements in multiple aspects of health span, resulting in significant delays in the appearance of pathology and frailty. The rationale for testing the drugs in combination as a single cocktail was based on the different but overlapping anti-aging targeting effects of each drug on processes of aging. C57BL/6 (B6) and HET3 4-way cross (H3) mice, 23 months of age, fed a diet containing the cocktail for three months showed significant strain and gender-dependent improvements in biological and physiological assessments and suppression of age-related lesions compared to mice fed individual drug or control diets.

The major biological changes in mice treated with the drug cocktail for three months were in body weight and percent body fat. Body weight was decreased by the cocktail in H3 males and females, but not in B6 mice of either gender. There were no decreases in body weight in either gender of either strain fed diets with individual drugs. Therefore, the combination of the three drugs had a bioloigical effect while each drug by itself did not. Interestingly, a decrease in fat mass occurred in both strains and both genders fed the cocktail diet suggesting H3 mice have different metabolic responses to the cocktail, especially since these mice increased their food intake compared to B6 mice. Blood glucose levels were not consistently affected by the cocktail diet, as mice fed control diet often had levels similar to mice fed the cocktail diet. Effect of the cocktail diet on fat mass was striking in both strains and was not gender specific, but appeared to be due to Acb and not Rap or Pba. One of the explanations for these phenotypic responses might be caused by the glucoregulatory control effect by Acb. Acb treatment selectively inhibits carbohydrate metabolism by allowing caloric compensation in food intake^24^. This selective inhibition promotes a similar effect as caloric restriction, which reduces calorie consumption provided by all macronutrients. Previous studies showed that Acb treatment led to lifespan extension principally in male mice^25^. Acb related metabolic pathways were similar in female and castrated males suggesting the sexual dimorphism effect was related to gonadal hormone differences. The gender bias was also reflected in this study where male mice showed more robust decrease in percentage body fat mass than female cohorts.

Several tests were conducted to measure physiological performance. B6 mice fed the cocktail diet showed improved cognitive performance, but H3 mice did not. Interestingly, Het3 mice showed less age-related cognitive impairment than B6 mice, but the cocktail had no effect in changing performance in this strain. When B6 mice were fed diets with each individual drug, only Rap had an effect in females, while all three of the drugs had an effect in males. Rab appeared to be the major contributor for the cocktail’s effect on suppressing cognitive impairment. Decreased neuronal activation and imparied cognitive peformance during aging occurs in both humans and rodents. Chronic mTOR attenuation by rapamycin has shown the benefits of restoring deficits in neurovascular couping response and cerebovascular dysfunction in aging rodent models^26^. Similar effects may be occurring in the current study.

Additional performance tests included a rotating rod and grip strength. B6 females fed the cocktail diet stayed on the rotating rod longer than the other cohorts. Interestingly, B6 males and H3 females on the cocktail diet did no better than respective mouse groups on the control diet. H3 males on the cocktail diet were underperformers staying on the rotating rod less than half the time of mice in the other cohorts. On the other hand, both strains and genders of mice fed the cocktail diet had increased grip strength compared to mice fed the control diet suggesting the cocktail was more effective in targeting pathways involved in muscle strength rather than coordination to stay on a rotating rod. In order to see if a partciular drug in the cocktail was the major contributor to effects seen in the performance tests, B6 mice were fed diets containing each drug, or the cocktail or control. No single drug consistently enhanced performance compared to the cocktail. However, female mice fed the Rap diet showed performance on the rotarod similar to mice fed the cocktail diet, while male and female mice fed the Acb diet were equal to cocktail treated mice in grip strength. The different results in the performance tests suggest that there may be selectivity regarding which component of the cocktail, and/or specific pathways, on the physiological outcomes we measured here.

In order to determine if a lower dose of the drugs in the cocktail might be effective over the three-month treatment period, B6 mice were fed a cocktail diet containing one-half the dose of each drug compared to the full dose cocktail diet and the control diet. Interestingly, the half-dose diet was just as effective as the full dose diet in decreasing body fat mass and preventing age-related cognitive impairment. In the physical performance tests, half-dose cocktail treated mice performed as well as mice treated with the full dose cocktail. However, the half-dose cocktail had no effect on reducing pathological lesiosns, suggesting again that there is some specificity in the outcomes beased on the dose of drugs used. Higher concentrations of the drugs were not tested, so it is not known whether a higher dose cocktail would have had a more comprehensive effect. However, the cocktail drug dose used in our study was adequate to show significant differences between mice treated with the various drug diets or control diet.

Age-related composite lesion scores can provide useful information as to how drug treatment can effectively target systemic organs^27^. Data from this study showed strain and gender differences in the composite lesion score of heart, lungs, liver and kidney in mice fed the cocktail diet. While B6 females and males and H3 females fed the cocktail diet had decreased lesion scores, age-related lesions in H3 males did not respond to the cocktail diet. This was an unexpected observation, which is difficult to explain. More insight might be gained by comparing the lesion scores of each individual organ in these mice. When individual drugs were compared to the cocktail in B6 mice, only the cocktail effectively decreased lesion scores in all four organs in both males and females suggesting the possibility of some type of synergistic activity among the three drugs that each individual drug was not able to provide. Another explanation would be that the wider targeting scope of the three drugs in combination for pathways of aging could be the difference in reaching the threshhold of positive effects.

In order to reveal possible molecular targets associated with changes in response to the cocktail diet, RNA message levels were detemined for p16, p21, TNFa, IL6, and IL1b in the kidneys from cocktail treated mice and control mice. The lower expression levels in cocktail treated mice were consistent with low expression levels in the kidneys from young untreated mice suggesting the cocktail delays aging partly by senolytic and anti-inflammatory effects.

In conclusion, the drug cocktail was generally more effective than each individual drug, and a half dose of the drug cocktail in the diet was overall less effective than the full dose cocktail based on biological, physiological and geropathological endpoints. The strain and gender differences partly agree with previous observations in lifespan studies in mice treated with Rap or Acb. Interestingly, previous studies also suggest that combining pro-longevity treatments may have beneficial effects on longevity and healthspan beyond that of each drug by themselves. For example, combining rapamycin and metformin extends the lifespan of HET3 mice at least as long as rapamycin alone and improves glucose metabolism Geropathology assessment showed that the individual Rap, Acb or Pba diets were no more effective than the control diet in decreasing composite lesion severity suggesting an interactive advantage of the three drugs adminstered together over the same time period. The lower expression levels of p16, p21, TNFa, IL6, and IL1b in the kidneys from cocktail treated mice compared to control treated mice were consistent with levels in the kidneys from young mice suggesting the cocktail delays aging partly by senolytic and anti-inflammatory effects, at least in the kidney.

## Acknowledgements

R56 AG058543, R01 AG057381, P30 AG013319, R01 AG057431

## Data availability

All data analyzed during this study are included in this published article.

## Disclosures

None of the authors disclose any competing interests.

## Notes

### Competing Interest Statement

The authors have declared no competing interest.

